# Beneficial microbes may have favoured the evolution of adaptive immunity

**DOI:** 10.64898/2026.01.29.702477

**Authors:** Lucas Mathieu, Thomas Pradeu, Richard Watson, Kevin Lala

## Abstract

Vertebrate adaptive immunity has long been considered uniquely advantageous for its capacity to recognise and remember a wide diversity of molecular targets, potentially conferring lasting defence from any threatening microbe. However, over the last two decades, results from comparative immunology have called into question the relative benefits of vertebrate adaptive immunity as a defence mechanism, raising the possibility of alternative evolutionary explanations. For instance, the hypothesis that managing the diverse and often beneficial host-microbiota associations characteristic of the vertebrate gut may have favoured the evolution of adaptive immunity. Here, we use individual-based simulations to investigate co-evolutionary interactions between hosts and their microbiota and the implication for the evolution of adaptive immunity. We focus on the capacity of adaptive immunity to produce and modify a diverse repertoire of immune receptors by using somatic variation and selective clonal reinforcement, in combination with a reinforcement bias from the innate immune system. Strikingly, we find that a high diversity of rapidly evolving parasites is insufficient to favour the evolution of adaptive immunity. Instead, our findings suggest that neutral and beneficial microbes can play a crucial role in weakening selection on innate immune defence against parasites, in favour of the exploitation of beneficial microbes. In turn, this facilitates the emergence of adaptive immunity as a mechanism reducing the risk associated with infrequent parasite infections while exploiting beneficial microbes. In addition, we show that adaptive immunity can subsequently alter selection affecting both innate immune defence and microbe evolution, such that its loss becomes increasingly unlikely with evolutionary time. Together, these results suggest that simultaneous interactions with diverse mutualists and parasites is a compelling selective explanation for the emergence and maintenance of adaptive immunity.

## 1 Introduction

Vertebrate adaptive immunity is generally defined as the capacity to generate lineages of cells expressing somatically-diversified receptors used in the binding, recognition and potential elimination of microbes [1]; and to re-mobilise the receptors acquired during the initial response to a pathogen (e.g., using long-lived cells) to accelerate subsequent responses to similar stimuli [2]. This process is called ‘adaptive’ because it allows hosts to generate receptors in response to novel immune challenges through stochastic and feedback-dependent phenotypic variation *within lifetime* [3] instead of relying on genetic variation and natural selection across generations. Although vertebrate immune systems vary largely, this core functional pattern is seemingly uniquely conserved across vertebrates but absent in invertebrates, and is thought to have evolved in ancient fish ∼500 Mya [4, 5]. However, generating and updating a reservoir of new receptors is presumably costly because stochastic and pre-emptive variation often produces significant amounts of unhelpful phenotypic variation. For instance in mice, a large proportion (up to 90%) of B lymphocyte precursors are eliminated to prevent auto-immune reactions [6]. In addition, among the staggering diversity of lymphocyte receptors produced within a vertebrate host’s lifespan, only a fraction is susceptible to react to specific antigens (e.g., a recent study found only ∼ 0.003% of human naive B cells were specific to the receptor-binding domain of SARS-CoV-2; [7]).

For several decades, the evolutionary trade-off imposed by generating costly and potentially ineffectual phenotypic variation has been explained by the presumed defensive benefit afforded by the flexibility offered by the under-determination of initial adaptive immune responses by host genomes and by the rapid secondary responses produced subsequently [1]. However, the idea that adaptive immunity is a markedly beneficial form of defence has recently received increased scrutiny [8], for at least three reasons. First, immune memory is broadly spread across invertebrates and plants and is even present in cells considered as part of the innate immune system in vertebrates [8, 9, 10], while the efficacy of long-term immune memory starkly varies across vertebrates [11]. Second, supposing that adaptive immunity is a superior system of defence raises the question of why a similar mechanism did not evolve in invertebrates. After all, vertebrates constitute most of the biomass and biodiversity on earth; why would they not evolve and benefit from this ‘superior’ defensive system, too [12]? Third, adaptive immune responses are often dependent on the initial recognition and capture of microbes by innate immune cells [13]. Together, these objections have motivated further hypotheses to explain why adaptive immunity evolved and was subsequently conserved across jawed and jawless vertebrates but not in invertebrates [14, 15, 16, 12]. Multiple authors have approached this question by arguing that constraints on evolvability, rather than selective explanations, are responsible for the absence of functional counterparts to adaptive immunity in invertebrates —besides immune memory, often called “trained immunity” in non-vertebrate systems [17]. For instance, by proposing that invertebrates lacked innovations such as the RAG transposons involved in somatic recombination [18, 19] or the available genomic complexity required for the regulatory control of adaptive immune responses [16]. In the following, we focus our analysis on a particular selective explanation, but we consider the influence of developmental and genetic constraints in our Discussion.

Here, we investigate the hypothesis that adaptive immunity evolved not solely for defence but rather to manage interactions between the host and a diverse microbiota. That is, it evolved to “combat pathogens while maintaining a tolerant relationship with symbiotic microorganisms” [20]. This argument was initially proposed by McFall-Ngai (2007) and further discussed by Lee and Mazmanian (2010) and others [23, 24] based on two principal motivations. First, extensive empirical evidence now points to the intricate, multi-faceted and often beneficial interactions between vertebrates (particularly mammals) and their gut microbiota, which are often mediated by adaptive immunity [25, 26, 27, 28]. Second, invertebrates generally form associations with fewer resident microbes than vertebrates [29]. Often, these associations involve a single specific microbial partner, with a few exceptionally diverse microbiota such as in termites and cockroaches [30], [31]. From the host’s perspective, matching the high diversity of gut microbes —–both beneficial and threatening–— constitutes a challenging evolutionary problem. Innate immunity is traditionally viewed as constituted of conserved immune pathways such as TLRs that target common microbe-associated molecular patterns (MAMPs), which may limit distinctions between harmful and beneficial microbes and impede the ability to foster a diverse and beneficial microbiota [32] (but see [33, 34]). By contrast, matching a diverse community of microbes with a somatically diversified repertoire of receptors may seem like a more parsimonious evolutionary solution [22]. However, recent discussions of a microbiota-driven adaptive immunity have primarily been motivated by contemporary immunity-microbiota interactions [35, 36, 31], which may not accurately reflect interactions across ancient evolutionary history. Crucially, a mechanistic account of how interactions with the microbiota could facilitate the emergence and subsequent conservation of an ancestral form of adaptive immunity is still lacking, to our knowledge. Below, we propose a minimal mechanistic model to evaluate this hypothesis.

We focus our analysis on a common functional pattern conserved in vertebrate immunity [37, 4] characterised by 1) the generation of somatic variation in receptor genes exclusively expressed by distinct clonal cell lineages, leading to a diverse anticipatory repertoire [38]; and 2) the differential multiplication of clonal receptor-expressing cell lineages mediated by affinity-dependent competition for antigen binding. Assuming that hosts can sample some of the microbes present in their gut, this is sufficient to allow the proportion of receptors with high binding affinity to follow the proportion of their corresponding antigens, potentially leading to highly specific immune responses [1, 39]. The precise mechanisms involved in the initial generation of receptor diversity and subsequent selection differ between jawed and jawless vertebrates [37, 40, 4, 5, 41, 16]. However, in both vertebrate lineages this process of somatic variation and selection generates two archetypes of receptor-expressing cell types (hereafter ‘lymphocytes’): a regulatory subset (i.e., VLRA in jawless fish and T lymphocytes in vertebrates) and a secretory subset (respectively VLRB in jawless fish and B cells). We focus our reasoning on the latter archetype, on account of its high conservation and direct role in regulating microbe-host interactions in the gut —sometimes independently from the intervention of the regulatory archetype in the case of mucosa-associated secretory lymphocytes [42, 43, 24, 44]. Admittedly, regulatory lymphocytes also play a key role in host-microbe interactions [45] and might have evolved hand-in-hand with, or shortly following, humoral immunity to reduce the risk of ‘self’-directed responses [46, 24]. For simplicity, here we assumed that regulatory control evolved as required by humoral immunity and we focused on interactions at the gut mucosal interface. We refer to this restricted model of humoral adaptive immunity as ‘somatically diversified immunity’ (henceforth SDI) to stress the importance of somatically diversified and selected receptors. We avoid the term “adaptive immunity” to prevent confusions between this minimal model and contemporary capacities of systemic adaptive immune systems, such as affinity maturation by hypermutation [39], long-term memory [47, 48] and complex regulatory control [49, 50, 51], which are not all conserved across vertebrates — see Rimer and colleagues (2014) for a comparative review in cold-blooded vertebrates.

To investigate whether interactions with the microbiota could have favoured the evolution of SDI, we consider an ancestral population of vertebrates with innate immunity co-evolving with a population of microbes. Using agent-based evolutionary simulations, we compare scenarios where the diversity of microbes and the evolutionary constraints affecting their evolution vary. In parallel, we assess the importance of host factors commonly assumed to influence host-microbe associations, including the shape of the dose-response function associating microbe density to host fitness. We show that SDI is favoured in hosts interacting with diverse populations of evolving microbes. However, a higher diversity of threatening microbes did not increase the frequency of SDI. Rather, we find that diverse and generally beneficial communities of microbes tend to weaken selection for innate immune defence, which in turn favours the emergence of SDI. We also examined the conditions in which SDI can subsequently be conserved across long evolutionary time, in an attempt to explain why SDI might be conserved in jawed and jawless vertebrates but absent in invertebrates. We show that SDI can disrupt selection on the innate immune system and stabilise the evolution of a higher diversity of beneficial microbes, both of which may contribute to its long-term conservation.

### Terminological considerations: mutualists, commensals, parasites and pathogens

Published works concerned with interactions between hosts and microbiota and their significance for adaptive immunity use a variety of terms to characterise microbes with beneficial (“partners” [21]; “symbionts” [52, 53, 54, 33]; “commensals” [32]) and harmful (“pathogen” [33, 44, 22, 32]; “pathobiont” [55, 56]; “parasites” [57]) effects on their hosts. But this categorical distinction is often artificial, with microbe ecology and evolution blurring the line between neutral, harmful and beneficial microbes [58, 59]. Generally, microbe “virulence” (i.e., direct damage to the host) is highly context-dependent [60] and can for example be plastically induced via quorum sensing [61] and other microbe-microbe interactions [62, 56]. Indeed, well-known human “pathogens”, such as *N. meningitidis, H. influenzae* and *E. Coli* are most commonly found as harmless commensals in hosts. Yet they can cause severe infections under opportune circumstances. Conversely, microbes classically viewed as beneficial, such as *Lactobacillus* and *Bacteroides*, can be harmful when multiplying in sensitive compartments of the host [63, 64]. Besides, ‘harm’ and ‘benefit’ are not restricted to direct effects on host tissues. When considering microbe-host interactions holistically, microbes consuming resources (‘public goods’) in the gut without producing nutrients or services that benefit the host, as well as those aiding the proliferation of virulent microbes [65, 56] also generate indirect costs to their host [66].

However, from a modelling perspective, capturing the intricacies of context-dependent microbial responses in an eco-evolutionary framework is largely impractical. Consequently, we adopt a coarser lens and consider that some microbes do have higher predispositions than others to detriment their host [67]. Evolutionary constraints on the availability of metabolic pathways result in convergent patterns of microbe adaptation across hosts producing similar habitats [68], and may lead to harm-prone groups. Thus, we simplify our approach by representing the general tendency of a group of microbe to cause harm or benefit considering their simulated evolutionary trajectory, rather than context-dependent harm (more on this in methods).

In addition, as shown by their pervasive use even in recent literature —a search in *The Web of Science* produces 1,782 results for “gut pathogen” for publications from 2024 alone— these labels still carry meaning in the scientific community. Accordingly, the opposition between stereotypically beneficial and harmful microbes is central in hypotheses linking the microbiota with the evolution of adaptive immunity [21] that were proposed by contrast with defence-centric metaphors of immunity as a response to “pathogens”. In the following, we adopt this coarse-view perspective to label groups of microbes that tend to detriment and benefit their hosts —respectively ‘parasites’ and ‘mutualists’, following similar existing computational analyses of microbes-host evolution [69, 70, 71]. Note, while viruses may play a role in these interactions, existing hypotheses have largely focused on microbes so far, and so does the present work.

## 2 Model summary

An overview of the model is presented in figure 1. We assumed that host fitness was entirely determined by the cumulative effect of beneficial and harmful microbes in their gut. Hosts influenced the microbial population in their gut by using either exclusively innate immunity or a combination of innate immunity and SDI. Innate immune reactions only depended on host genotypes and on the perceived harm caused by the microbiota, the latter mimicking inflammatory responses. Innate immunity was environmentally responsive but deterministic, such that two host clones with the same microbiota would produce the same innate immune response. By contrast, SDI was based on the stochastic population dynamics of multiple lymphocyte lineages within each host, which were selectively reinforced by antigens sampled in the gut. This antigen-mediated feedback was itself biased by the innate immune system. Instead of the common assumption that the principal constraint on immunity is the cost of auto-immune reactions [24, 72], we considered that implicit costs of immunity would arise from immune reactions targeting ‘mutualists’ and from insufficient immune responses to ‘parasites’ (see box above for terminology). Thus, we did not assume intrinsic costs of either form of immunity. Hosts reproduced asexually in partially overlapping generations with fitness-proportionate selection. Fitness was updated cumulatively for up to 100 time-steps per host. We tested multiple dose-response functions associating the density of each microbe species within the gut to its contribution to the host’s fitness. In the following, unless specified otherwise beneficial effects are modelled with a logarithmic response function whereas harmful effects are modelled with an exponential response. This aimed to reflect the potential saturating effect of nutritional intake in mutualistic associations, by contrast with common density-threshold effects associated with parasites that often only produce harmful effects at sufficiently high densities [61]. Each reproductive event, host genotypes could mutate to modify their innate immune response to a subset of microbes. This represented an implicit set of innate receptors and associated responses pathways, which we assumed to evolve hand-in-hand. After co-evolutionary dynamics between hosts and microbes stabilised (usually 3*e*4 time-steps; see figure 4) we let SDI invade populations with a low mutation probability. For simplicity, we only considered antagonistic effects of host immunity towards microbes in the gut, assuming positive effects —such as the provision of nutrients like mucin [73]— to be implicit and shared across all microbes.

**Figure 1.**
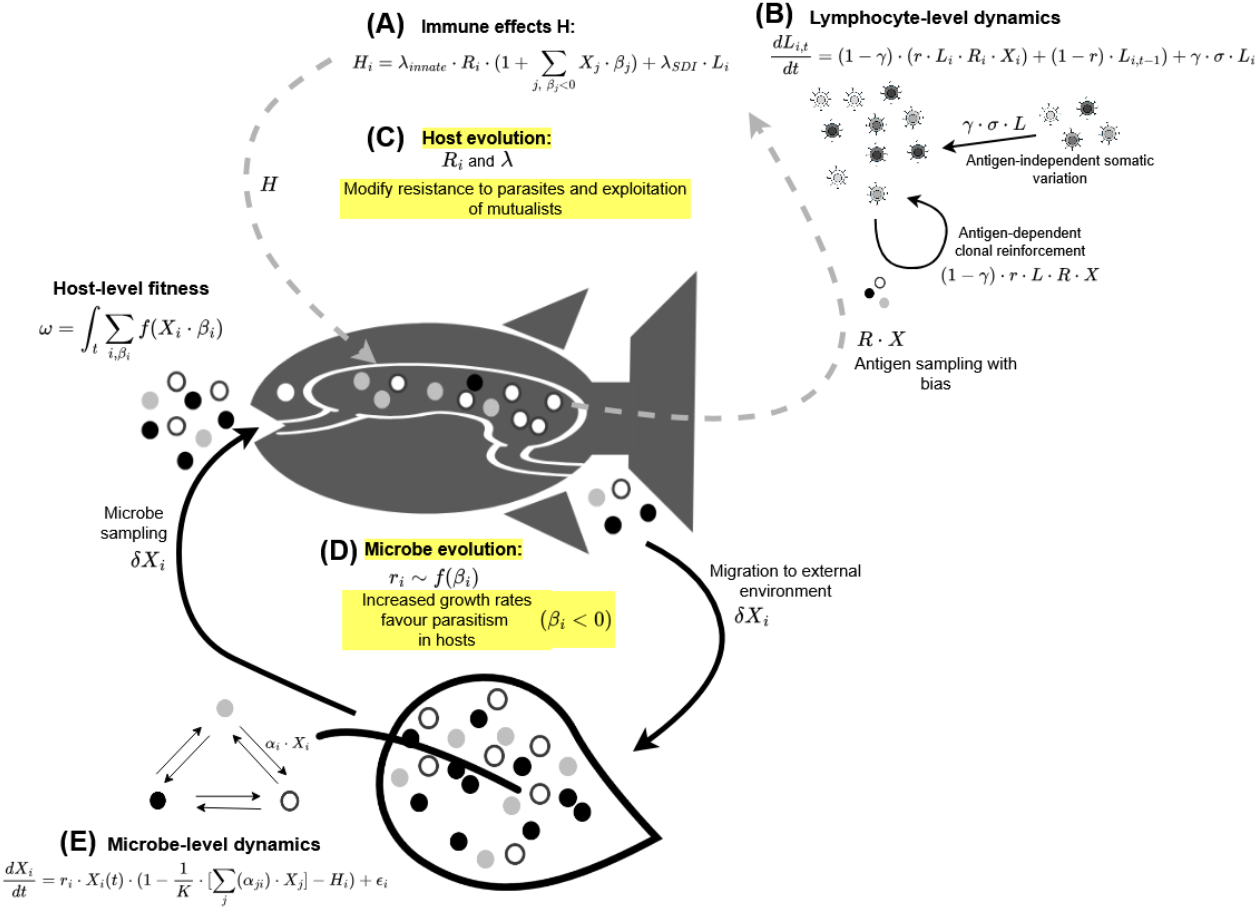
Overview of the model. We consider a population of hosts representing vertebrate ancestors, whose fitness ω only depends on microbe-host effects β weighted by microbe densities *X* and a dose-response function *f*. Microbes are sampled from the environment following a hurdle model with a Bernoulli trial of probability ϵ and normally-distributed amplitude. E: Microbe dynamics depend on their growth rate *r*_*i*_, linear interaction terms α_*ij*_ with other microbes and host immune effects *H*_*i*_ (null outside of hosts). A: Hosts influence the population of microbes in their gut using an immune response *H* determined by an innate component and a facultative SDI component. Innate immune responses are proportional to the perceived weighted harm inflicted by microbes β_*i*_*X*_*i*_, β_*i*_ < 0. B: SDI immune responses are formed by a variation-selection process in a population of lymphocytes *L*_*i*_ targeting each microbe species *i*. Variation is produced by stochastic and antigen-independent influx of lymphocytes γ · σ · *L*, following a Gaussian hurdle model of probability γ and amplitude σ. Lymphocyte lineages are selectively reinforced based on antigens sampled by the innate immune system in proportion to the gut content *R* · *X*. C: Host can evolve by modifying their innate repertoire *R*, by modifying the overall intensity of their immune responses λ and by adopting or losing SDI, which is controlled by a discrete gene *µ* taking values 0 and 0.5. D: Microbes evolve by modifying β_*i*_ towards parasitism (β_*i*_ < 0) or mutualism (β_*i*_ > 0), which is driven by a positive relationship between parasitism and growth rate within hosts and a negative or neutral relationship between parasitism and growth rate in the external environment.

We let microbes evolve within hosts and in the external environment, such that parasites contributing negatively to host fitness have a higher growth/death rate *r* within hosts (figure 1, D). Microbes that are targeted by the immune system of their host do not benefit from higher growth/death rates, because the immune contribution makes their net growth negative (figure 1, E). Therefore, the host’s immune systems can subject parasites to random drift, displacing them on the mutualism-parasitism spectrum and countering selection for parasitism. However, to maximise the benefits received from exploiting mutualists the hosts must avoid suppressing them with immune responses, which lets mutualists evolve towards parasitism. This produces a difficult evolutionary challenge for hosts because previously beneficial microbes can gradually become threats, which may prevent the maintenance of long-term mutualism [70]. Hosts evolved by modifying their innate response to a given microbe species *i* through immune genes *R*_*i*_, which also biased responses from SDI; and by modifying to the relative strength of innate and SDI immune responses in λ_*innate*_ and λ_*SDI*_ (A and C). To constrain and stabilise microbe evolution, we also let microbes evolve in the external environment where microbe-microbe interactions are the principal driver of selection, which constrains and stabilises host-microbe co-evolution. In short, host innate immune systems must keep up with microbes such that current threats are suppressed while beneficial microbes are left alone. In these conditions, the inherently stochastic nature of SDI produces a large direct cost by potentially targeting mutualists —while also delaying immune responses to parasites. In addition, the developmental noise generated by SDI produces a second indirect cost by disrupting the inheritance of fit innate immune systems, hindering their evolution. Conversely, when the innate immune system is error-prone the stochastic (albeit delayed) density-dependent responses of SDI may become beneficial by limiting the risk posed by abundant parasites that are not sufficiently suppressed by host innate repertoires.

We investigated how the above trade-off may respond to the influence of four principal variables. 1) The number of microbe species in the environment, *n*. Each ‘species’ represented by a) a marker *i* serving as a target for the hosts’ immune system; b) a microbe-host interaction term β_*i*_ characterising whether this species is beneficial or detrimental to the hosts; and c) a matrix of microbe-microbe interactions α_*i,j*_ representing competition and cooperation between microbe species. For simplicity, we assumed that microbe-microbe interactions did not change during our simulations because these were not the primary focus of this work. We let microbes evolve by changing the sign and amplitude of their effect β_*i*_ on hosts, thus varying on the mutualism-parasitism spectrum. We made the typical assumption that parasites multiplied faster than mutualists within hosts [74, 52, 75, 70], which created a selective pressure for parasitism. Conversely, parasitism did not necessarily increase growth rate in the external environment. 2) Additionally, we varied the relative advantage of parasites and mutualists in the external environment to indirectly control the proportion of beneficial and harmful microbes by constraining their evolution. 3) We also varied the relative intensity of microbe-microbe competition to explore the potential role of competitive exclusion. Finally, 4) we investigated how the dose-response function of the host to the abundance of beneficial microbes influenced the evolution of SDI. A detailed account of the model and parameter choices is given in section Materials and Methods.

## 3 Results

### 3.1 A high diversity of beneficial microbes favours the evolution of SDI

As described in the model summary, SDI is costly because it produces stochastic variation in immune responses, whereas innate immunity can deterministically target parasites but if selection on host genomes can ‘keep up’ with microbe evolution. However, when the evolution of innate immune systems is outpaced by microbial evolution SDI might be beneficial by limiting the threat posed by parasites that would escape the host’s innate immune responses. As a consequence of this trade-off, we find that SDI only evolves in restricted conditions (figure 2). First, SDI only evolves when the diversity of microbe species in the environment and within hosts is high (>60 species). This supports the general intuition behind the ‘care for the microbiota’ hypothesis: when the number of microbe species commonly interacting with hosts is low, selection on the innate immune system can rapidly follow changes in microbe populations. Therefore, the costs potentially posed by SDI are high, making innate immunity preferable. Conversely, when the number of microbe species —each potentially capable of evolutionary switches between mutualism and parasitism— is high, the defence-focused paradigm of adaptive immunity may predict that innate immunity should be overwhelmed, making SDI a better alternative. Our results partially support this view, but with a significant caveat. Indeed, we find that even when microbe diversity is high SDI is rarely favoured when parasites are dominant (figure 2), panel A). In such conditions, hosts with innate immunity tended to tolerate very few microbes while suppressing the vast majority of others, effectively simplifying the evolutionary challenge posed by the larger diversity of microbes (see figure 4, panel D: microbe population evenness is markedly lower without SDI, indicating that few microbe species are dominant while most others are virtually absent). By contrast, we find that SDI is favoured when the proportion of mutualists is high, particularly when microbe-microbe competition is intense. This suggests that the benefits of SDI are more pronounced when the density of any given microbe within host is kept low by competitive interference, such that the risk of harm from any given parasites is lower. In short, when parasites pose a greater risk, they generate strong selection on the host’s innate immune system. This forces hosts to ‘keep up’, at the cost of collateral immune responses targeting mutualists. In these conditions, SDI is strictly detrimental. Conversely, when parasites are comparatively less threatening they produce weaker selection than mutualists on the immune system of hosts. This leads to an imbalanced trade-off between defence and tolerance favouring SDI as a risk-mitigating strategy in hosts exploiting diverse mutualists.

**Figure 2.**
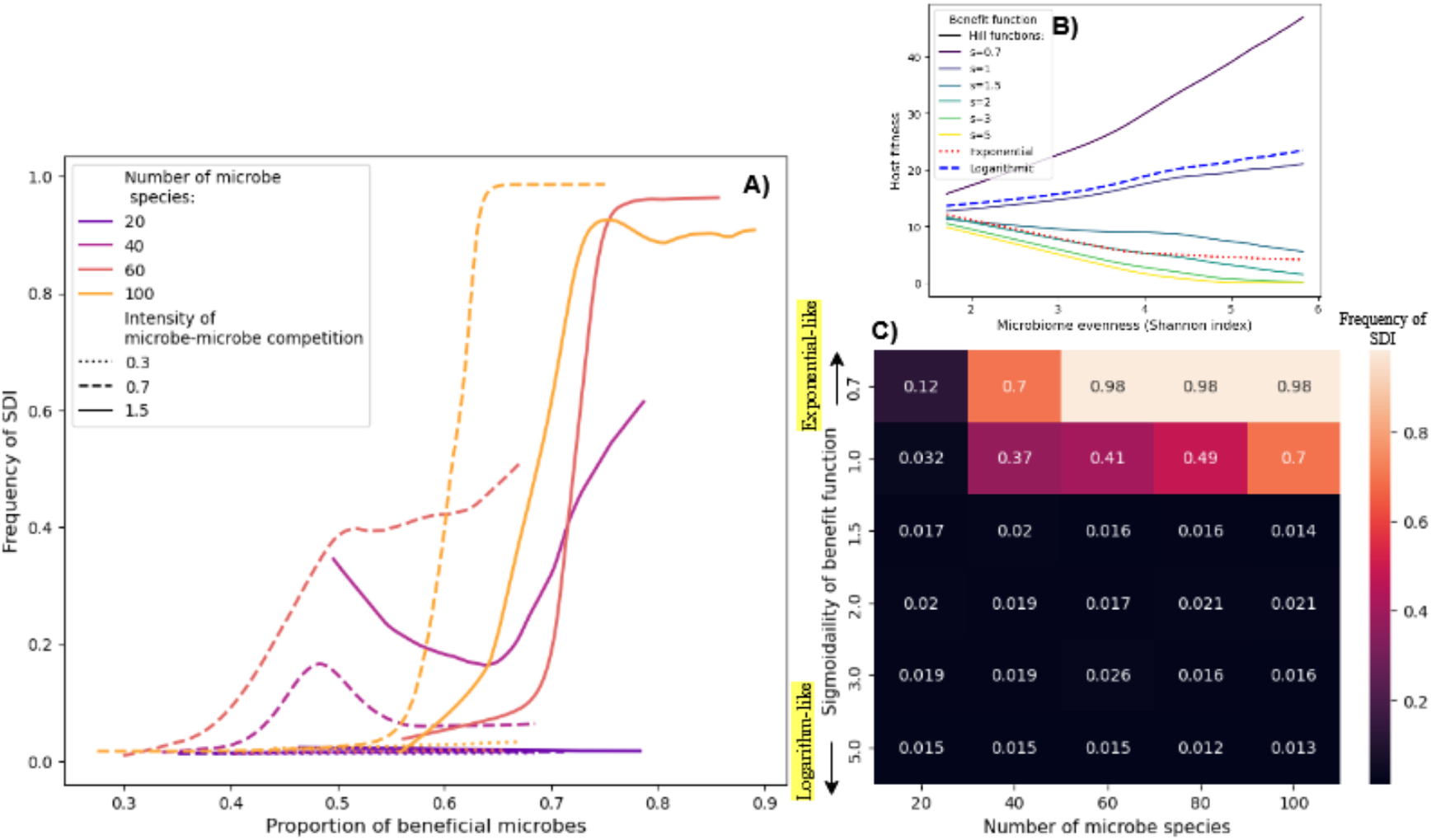
Somatically diversified immunity (SDI) is only favoured by diverse and predominantly beneficial microbial communities. A: We indirectly control the proportion of beneficial microbes by modifying the sign and amplitude of the trade-off incurred by microbes in the external environment: and observe the frequency of SDI after evolutionary dynamics have stabilised. We find that SDI is only likely to evolve when the majority of microbes (at least 55%) are beneficial. In addition, the requirement of a predominantly beneficial microbiota was alleviated by increased microbe-microbe competition (full lines), indicating an effect of competitive exclusion. B: We examine the influence of the benefits function of hosts by modifying the parameter *s* (‘Sigmoid factor’) controlling the curvature of a dose-response function defined as a Hill function. When *s* is low, the weighted beneficial effects of microbes increase very rapidly with microbe density but plateau rapidly, such that hosts benefit more from exploiting a high diversity of microbes. When *s* is high, weighted positive effects from microbes follow an exponential shape such that hosts only receive large benefits from microbes present at high densities. This relationship is shown in C by the interaction between microbe community evenness and host fitness with respect to the benefit function. Hosts with a concave (*s* <= 1 and Logarithmic) response functions benefit from even (high Shannon index) microbiota, whereas hosts with a convex (*s* > 1 and Exponential) benefit function are favoured by uneven microbiota. We find that SDI only evolves when microbe diversity is high and hosts benefit more from exploiting many mutualists rather than one or few dominant microbes in an uneven community (top right corner of the heatmap).

### 3.2 SDI only evolves when hosts benefit from exploiting multiple microbe species simultaneously

Hosts generally do not linearly benefit from an increase in the abundance of a given mutualist species, nor do they proportionately benefit from an increase in microbe species richness. Depending on the number of microbiome ecological services that the host relies on, the loss of a single microbe species may be dramatic or fully inconsequential [76]. Borrowing the metaphor of “rivet-redundancy” *vs* “immediate catastrophe” [77] in ecological resilience studies, we analysed the effect of the sensitivity of hosts to microbiome diversity (e.g., from a nutritional perspective) on the evolution of SDI (figure 2, panel B and C). There was always a saturation threshold in the dose-response function. However, this saturation could either be reached rapidly, favouring hosts with an evenly distributed microbiota; or slowly and with few benefits for hosts at low microbe abundances, making microbiota with one or few dominant mutualists preferable. We find that SDI only evolved when host fitness was maximised by more even microbiota. Specifically, SDI was favoured by convex benefit functions (either logarithmic or logistic with a low curvature rate *s* <= 1) but never evolved when weighted microbe-host effects followed a concave (either exponential, logistic with *s* > 1 or step-like) shape with low benefits at low densities and higher benefits at higher densities.

This can be interpreted as follows. On the one hand, SDI produces stochastic and density-dependent responses that target abundant microbes, which hinders the ability of hosts to foster any particular mutualist species in high abundances. On the other hand, these same characteristics are ideal to maintain a balanced population of microbes within the host. Consequently, SDI evolved when hosts benefited more from an evenly distributed community of diverse mutualists.

### 3.3 Disrupted selection on innate immunity favours SDI and is self-sustaining across long evolutionary time

Results from the previous section show that the conditions favouring SDI are limited to environments with a high diversity of predominantly beneficial microbe species, where microbe-microbe competition is high (fig 2, panel A) and where hosts benefit more from an evenly distributed community of diverse microbes than from a few dominant mutualists (panel C). These conditions are unlikely to apply across all vertebrate lifestyles and across their entire evolution, even if we assumed that such an environment arose at some point in vertebrate ancestry; unless the fixation of SDI in vertebrate ancestors produced and maintained at least some of the conditions that favour its own maintenance.

#### 3.3.1 SDI disrupts selection on the innate immune system

In figure 4 (panel B), we show that the presence of SDI disturbs selection on the innate immune system, preventing the evolution of receptors that suppress parasites while ignoring mutualists. This is particularly marked when the evolutionary task is made more demanding by a high proportion of beneficial microbes, such that mutations to the innate immune system that improve defence against a parasite are likely to also increase the suppression of multiple beneficial microbes (fig. 3, right-most panels). In other words, we find that with time, the innate immune system becomes increasingly poorer at suppressing parasites while ignoring mutualists in populations of hosts where SDI has been fixated. Consequently, a loss of SDI and reversion to relying exclusively on a deprecated innate immune system becomes increasingly unlikely as the associated innate immune system becomes gradually incapable of functioning on its own. This is shown in figure (3, panel A) by a plateau near 0 in the probability of a decrease in the frequency of SDI as its cumulative frequency (i.e., the time spent after SDI has been fixed in a population of hosts) increases. This is particularly true for higher numbers of microbe species (*n* >= 80). This shows that mutational noise on the trait ‘presence/absence of SDI’ is largely insufficient to remove SDI from populations in which it has evolved.

**Figure 3.**
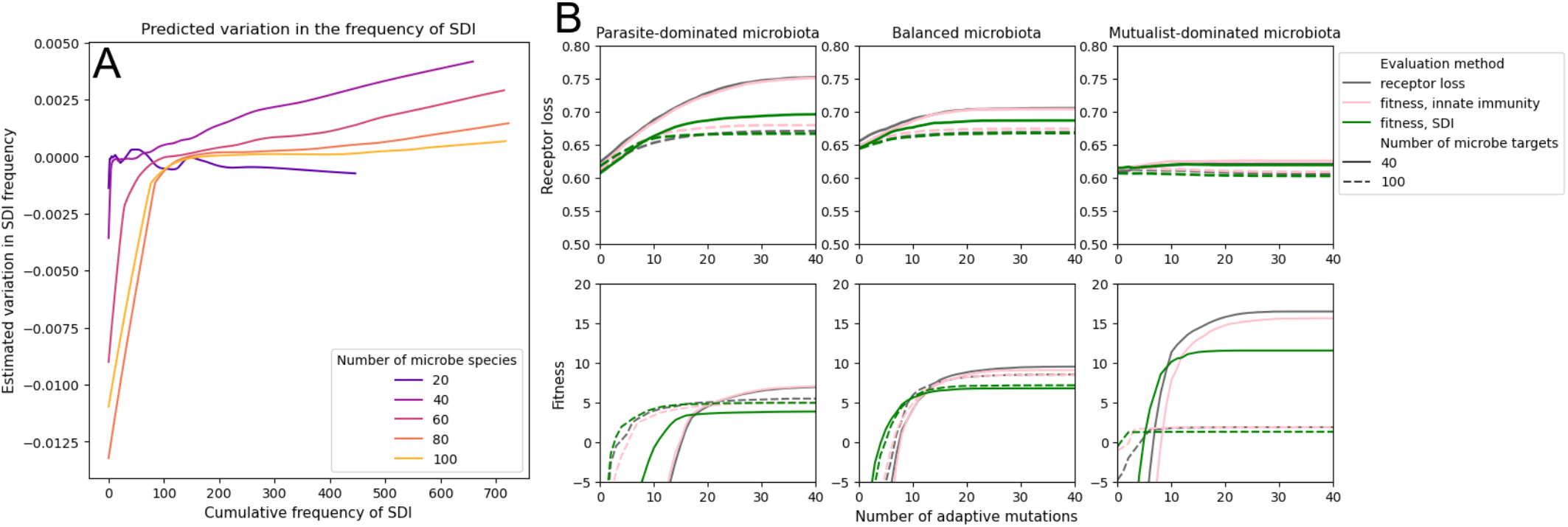
The persistence of SDI is self-stabilising. A: When microbe diversity is sufficiently high in the host gut microbiota, the probability of loss of SDI plateaus near 0 after a few generations. B: The presence of SDI hinders the evolution of innate immune receptor repertoires. We assess the performance of innate immune repertoires with a simple criterion ‘Receptor loss’: innate genes *R*_*i*_ must target parasites while avoiding mutualists. This criterion is illustrated in panel B, top row in shorter simulations where microbes did not evolve. Innate repertoires incrementally mutated using this criterion are indistinguishable from the innate repertoires that would naturally be selected using a fitness-based evaluation (grey and pink lines, respectively). Conversely, the presence of SDI disturbs selection on innate repertoires, delaying and lowering the plateau of performance both expressed as fitness and as ‘receptor loss’.

#### 3.3.2 SDI promotes the evolution of a stable and beneficial microbiota

In addition to affecting the evolution of the innate immune system, the presence of SDI markedly modifies the evolutionary trajectories of the associated microbiota in our simulations (figure 4, panels D and E). Despite not increasing fitness (and even contributing to a marked drop in fitness, panel B), the evolution of SDI markedly affects the associated microbiota (green line at *t* = 3*e*4, panels D, E and F). Specifically, microbe populations interacting with hosts relying on SDI become markedly more diverse (species richness, panel D), with increased differences between hosts (panel E). However, this is not associated with a marked change in the tendency of microbes to mutate from mutualism to parasitism and vice-versa (panels F and G).

**Figure 4.**
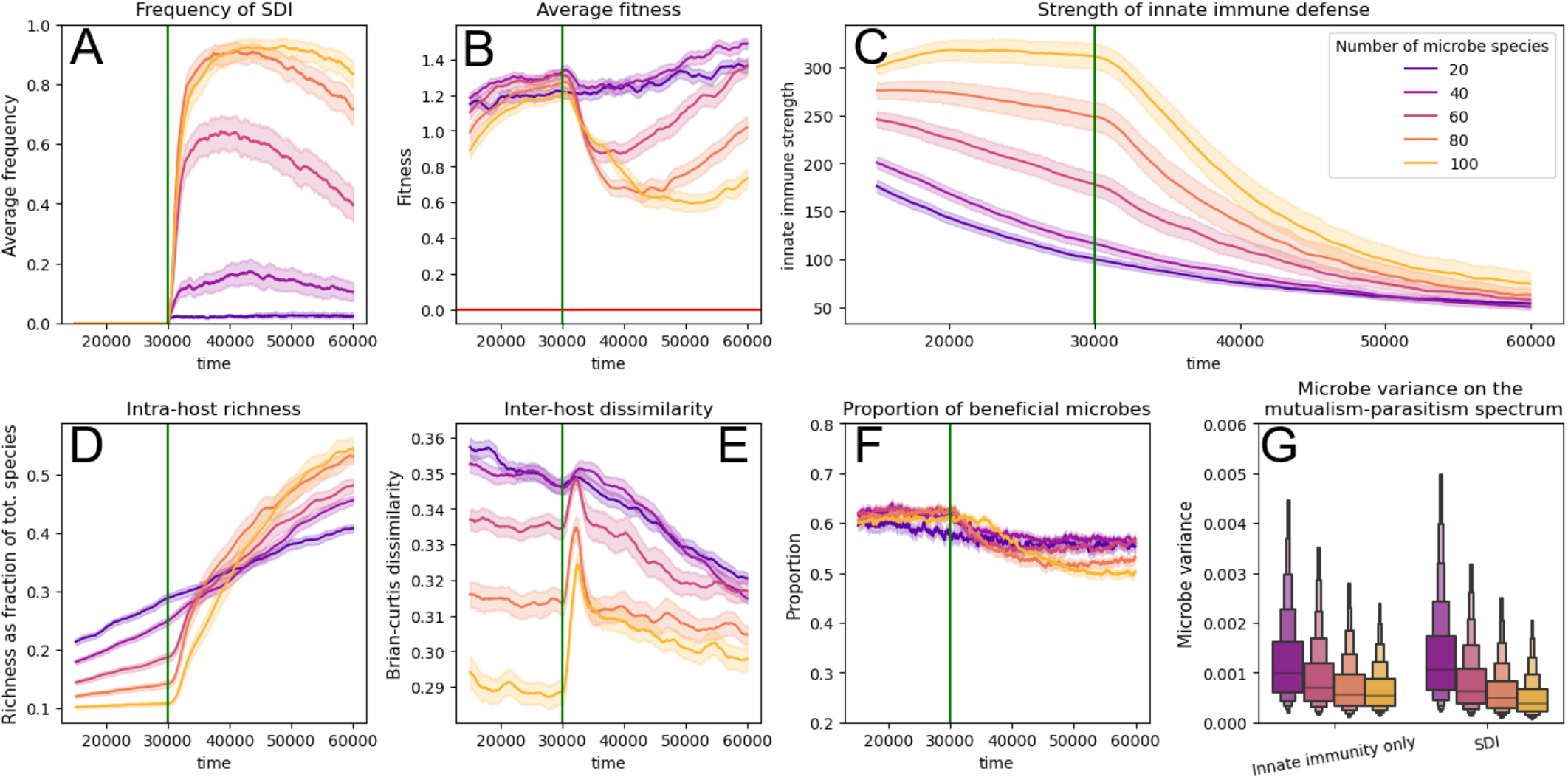
Co-evolutionary interactions between SDI and the microbiota. Microbes were subjected to a slight trade-off in growth rate between host and external environment (τ = 0.02). Microbe-microbe interactions were competitive on average (α_*i,j*_ ∼ 𝒩 (0.7, 0.3)). After evolutionary dynamics between hosts initially using exclusively innate immunity and their microbiota have stabilised, SDI was allowed to invade populations by clonal mutation (green vertical line). A: SDI only evolves when the diversity of microbes (i.e., total number of species that can be encountered) is high. B: The fixation of SDI is marked by an abrupt decrease in fitness, that we attribute to its immediate effect on microbe populations dynamics and on innate immunity. C: The average strength of innate immune defence rapidly decreases in populations where SDI has evolved (number of microbe species *n* >= 60). D: SDI markedly increases microbe species richness (average number of microbe species with non-zero abundance within a host, relative to to *n*) within hosts. E: The evolution of SDI is also marked by a sharp increase in microbiota dissimilarity between hosts. F and G: The evolution of SDI has little effect on the overall tendency of microbes to vary on the mutualism-parasitism spectrum at the meta-population level, despite the marked ecological changes within and between hosts (D and E).

In addition to these changes, the evolution of SDI is associated with a rapid decrease in the strength of innate immune defences (fig. 4, panel C). This indicates that, irrespective of which microbes are targeted, suppressive responses from the innate immune system are on average weaker in the presence of SDI. Given that immune suppression from SDI depends on bias from the innate immune system that samples target microbe species from the gut and on the abundance of these targets, this shows a systemic shift from a focus on immune defence to a focus on tolerance in hosts with SDI. In hosts populations with a highly diverse microbiota (*n* >= 80) the decrease in the use of innate immunity to suppress microbes —following the evolution of SDI— is most apparent (panel C when *t* > 4*e*4). Together with the associated high proportion of beneficial microbes (fig. 4, panel F) and the requirement of high microbe-microbe competition for SDI to evolve (fig. 2), this indicates that competitive exclusion favours the evolution of SDI by gradually decreasing the need for a highly performant and defence-focused immune system, in favour of an immune system capable of tolerating many microbe species while infrequently reacting to potential threats.

## 4 Discussion

We considered a minimal model of adaptive immunity with limited affinity maturation and recognition specificity (such that adaptive immune cells and innate receptors could in principle be equally specific), without long-term memory and restricted to mucosal and humoral interactions at the gut interface. Our results suggest that even with such restrictions, the selective pressure generated by interactions with the microbiota could induce the evolution of an ancestral form of adaptive immunity, under the following conditions. First, hosts must interact closely with their microbiota, and in such a way that they benefit from a diverse microbial community rather than a single or few symbionts as observed in contemporary invertebrates [78]. Second, the microbiota must be highly diverse and in particular, must contain a high diversity of mutualists. Third, interactions between microbes must be prevalently competitive, allowing beneficial and neutral microbes to protect hosts from parasites in a form of “co-immunity” [27]. Under these conditions, disrupted selection on innate immunity —caused in part by competitive exclusion from beneficial microbes— weakens the defensive function of the innate system. This opens the way for the evolution of a noisy (and therefore costly) form of immunity relying on somatic variation and feedback-mediated reinforcement to suppress dominant microbes, reducing the risk of parasite infection while preserving diversity within the gut community. However unlikely these initial conditions may seem, we find that they could be self-preserving. We find that hosts interacting closely with their microbiota may favour the preservation of a high proportion of mutualists, either with innate immunity at low microbe diversity or with SDI when microbes are highly diverse. Further, our results suggest that adaptive immunity may maintain a high effective microbe diversity within hosts [79] and disrupt positive selection on innate immunity, making a return to an innate-only immune system improbable.

### 4.1 A scenario for the evolutionary exception of adaptive immunity

#### 4.4.1 Constraints on invertebrate genomes

To explain the absence of adaptive immunity in invertebrates, multiple authors suggest that evolutionary constraints, rather than selective pressures, facilitated the evolution adaptive immunity in vertebrates while preventing its apparition in invertebrates. Two notable events in vertebrate evolution are often mentioned in such explanations. First, the appearance of RAG-based recombination along with the cellular machinery necessary to generate and selectively reinforce somatically diversified receptor cell lineages has long been considered as an evolutionary “big bang” [19, 16]; that is, an extraordinarily rare event that conditioned the evolution of adaptive immunity. But this view has been challenged by recent evidence of other (convergent) mechanisms of somatic diversification of immune receptors in jawless vertebrates [80, 41] and even in some invertebrates [81, 82]. Similarly, the principles of cell-cell competition required for clonal selection may be more common than previously thought [83, 84]. Conversely, non-vertebrate chordates such as Cephalochordates possess RAG and genes analogues of the vertebrate Major Histocompatibility Complex (MHC) commonly associated with adaptive immunity, but lack functional equivalents of adaptive immunity [85]. Kasahara and colleagues (2010) suggest that a more compelling origin for the emergence of adaptive immunity may be the event of whole-genome duplication (followed by a second duplication in Gnathostomes) separating vertebrates from invertebrate chordates, where adaptive immunity is absent. The authors argued that the resulting increase in genome sizes may have facilitated the emergence of the genomic complexity required to regulate an adaptive immune system [86, 16]. However, this argument suffers from a flaw: in principle, genome-wide duplications could have accelerated the evolution of new innate receptors and response pathways and increased the immune genetic diversity of vertebrates instead of giving rise to adaptive immunity [87, 81]. Unless the ancestral trait of adaptive immunity —for instance, RAG-based somatic variation— produced a very marked fitness benefit, there is no reason to believe that an increase in genome size and genetic diversity would favour adaptive immunity over improved innate immunity. Our results may suggest a solution to this conundrum. We find that interactions with a prevalently mutualistic microbiota could result in a mismatch between innate immunity and microbe populations (figure 3, panel B; the pink and grey curves differ most when mutualists are dominant and microbe diversity is high). This mismatch primarily results from asymmetric selection for tolerance of mutualists over defence from parasites, rather than from constraints on genomes that would hinder the evolution of immune systems. Consequently, a sudden increase in the availability of genetic diversity and novelty though whole-genome duplication may have not improved the innate immune system of hosts, but could have favoured the appearance of SDI instead.

#### 4.1.2 Vertebrate and invertebrate microbial environment: same but different?

Arguably, ancestral vertebrates and invertebrates likely shared a similar environment; at the very least, there must have been overlaps between the ecological niches of some ancestral invertebrates and some vertebrates [88]. Despite this, differences in the way vertebrate and invertebrate ancestors experience external stimuli and mutational novelty could have induced distinct selective pressures, even in a shared environment. Accordingly, multiple specificities of the vertebrate body plan [89, 90, 12] and metabolism [57, 91] have been proposed as explanations for the evolution of adaptive immunity in vertebrates. For instance, Nakashima and colleagues (2018) remark that unlike in many invertebrates, for which the gut epithelia is often reinforced (e.g., by chitin in Ecdysozoa) and respiratory and digestive tracts are separated, the vertebrate digestive tract is formed of a single layer of cells and joint with the respiratory tract at the pharynx. This may have greatly increased the sensitivity of vertebrate ancestors to the composition of their gut microbiota —and to specific types of parasites such as ‘helminths’ [92]— while also facilitating the evolution of an immune system relying on the indirect monitoring and control of microbial population by exchanges through a thin epithelium at the pharyngeal and gut interface [42]. Our results offer potential targets to study how such differences in biological systems might influence the response to selection in immunity. For example, the relative permeability of the gut epithelia in vertebrates may constrain the dose-response curve linking mutualist abundance and host benefits, biasing hosts to favour a diverse and even community of microbes, or vice-versa.

Members of the Cephalochordates sub-phylum may be promising animal model candidates for such research. These basal Chordates possess many potential precursors to the emergence of a minimal adaptive immune system in vertebrate ancestors, including a single-layer epithelium, shared nutritional and respiratory function in the pharyngeal cavity, evidence of active phagocytosis, immune effector secretion in gut epithelial cells, as well as analogues of the MHC and RAG genes considered as hallmarks of vertebrate adaptive immunity (see Yuan and colleagues (2015) for a detailed account in Amphioxus). Yet, no evidence of an immune system based on recombination and clonal selection has been found. Whether Cephalochordate ancestors lost a primitive form of adaptive immunity or were subjected to different selective pressures from vertebrates has yet to be determined, to our knowledge.

#### 4.1.3 Explaining the long-term conservation of adaptive immunity

Proposing that adaptive immunity may not have evolved as a superior form of defence [82] raises a secondary question. If adaptive immunity is not widely ‘adaptive’, how can we explain its conservation across vertebrate phylogeny? Ward and Rosenthal (2014) attribute the impoverished state of innate immune repertoires in vertebrates [94] relative to invertebrates to disrupted selection caused by adaptive immunity, with a proposed shift of innate receptor function from recognition to regulation. We propose a similar selectively neutral explanation, borrowing from the framework of “constructive neutral evolution” [95] and “evolutionary ratchet” [96]. Our results suggest that the exploitation of a diversity of beneficial microbes could have canalised the appearance of adaptive immunity by disrupting selection on innate immune defence, paving the way for a functionally redundant and genetically under-determined system in the form of SDI. Subsequently, both the presence of adaptive immunity and of protective symbionts may have further weakened selection on the innate immune system and increased the reliance of hosts on adaptive immunity. In our simulations, this is shown by the decrease in the average strength of innate immune responses (figure 4, panel C) and the difficulty of selecting adaptive mutations in innate immunity in hosts with SDI (figure 3, panel B, green vs pink and grey lines). Crucially, such selectively neutral explanations are potentially robust to temporal and geographical environmental changes that would modify selection on a given trait. In a recent review, Hammer (2024) suggested that some hosts may be “addicted” to their microbiota across evolutionary time-frames. Such an effect has been experimentally demonstrated in *D. melanogaster*, where the presence of protective symbionts has been linked with weakened selection on immune defence [98]. We suggest that such interdependencies may have occurred broadly in vertebrate evolution and could explain the conservation of both adaptive immunity and host-microbiota interactions, even in cases where the evolutionary benefits are apparently unclear.

### 4.2 Immune systems as constructors of their own selection

The idea that hosts and parasites co-evolve is not novel [99]. However, here we considered microbes not as abstract antagonists in an evolutionary arms race with their hosts, but rather as dynamic components of an ecological community. Recent evidence indeed suggests that eco-evolutionary feedbacks may play a crucial role in co-evolution [100], for instance by imposing both ecological and evolutionary sweeps that produce marked phenotypic change in members of the microbiota [101] and limit genetic diversity, increasing the influence of drift [102]. As such, hosts can influence the evolution of microbes not only by imposing specific selective pressures within their gut but also by controlling the demography of microbial populations. The vertical [103, 70] and horizontal [104] transmission of microbes have been shown to respectively facilitate and disturb these demographic-level forms of control. Accordingly, our results suggest that when hosts exploit microbes well-adapted to the external environment, the bilateral exchange of microbes between hosts and the environment may itself be utilised to control microbe evolution, similarly to the coordinated recruitment and dismissal of microbes in the bobtail squid [105]. Thus, even with small-scale microbe migrations from hosts to the environment, the effect of each host’s immunity may alter the distribution of microbes not only within hosts but also in the external environment, even when it contains far more microbes than each individual host (figure 4, panel E). Accordingly, microbes originating from the vertebrate gut have been isolated in the environment surrounding hosts, including on artificial surfaces [106] and in soils contaminated with faeces [107]. Together, this forms a picture of host-microbiota interactions as a form of reciprocal niche construction, where hosts and microbes may modify their respective environments by disrupting and stabilising the network of interactions composing the microbial community [108, 109].

Our understanding of the interactions between residents of the gut is generally limited to studies in humans and mice relying on temporal and geographical -omics data to infer interactions from non-random patterns of distributions [110]; and the building of intricate knowledge-driven models to complete these largely correlation-based methods [111, 112]. However, the integration of various external environments and life-cycles into our view of microbial ecology is still lacking [113] despite recent progress in our understanding of microbiota in non-classic model animals [114] and free-living microbiomes [115]. Recent experiments have demonstrated the potential interest of controlling the demography of the microbiota, showing that k-selected microbiota promoted better health than r-selected microbiota in fish [116]. However, the ecological dynamics of microbes outside of hosts may be just as important for microbe-host co-evolution as within-host dynamics, as suggested by the importance of the evolutionary trade-off affecting selection of parasitism within and outside of hosts in our model.

### 4.3 Model limits and future perspectives

To keep the model tractable, it only features a limited representation of adaptive immunity. ‘SDI’ lacks both long-term memory and the capacity for auto-immune reactions, which are both significant determinants of the impact of adaptive immunity on host fitness [8]. In addition, many functional properties of modern adaptive systems —such as local regulatory control, hypermutation and poly-reactivity— are absent from our model. We also confined our analysis to interactions between the host and the gut microbiota within the gut (or the pharyngeal lumen; see below). This focus on a minimal representation has multiple advantages. First, as stated in the introduction we isolated characteristics of the adaptive immune system that are conserved across most vertebrates —including modern fish [11, 117, 37, 118]— in order to keep our results generalisable and applicable to hypothetical vertebrate ancestors where primitive forms of adaptive immunity emerged. This approach offers a second benefit. Assuming that adaptive immunity emerged as a form of feedback-mediated mucosal immunity vastly reduces the requirements for the regulatory control necessary to prevent auto-immune reactions towards diversified cell types in systemic immunity. In our opinion, this simplified and highly conserved functional view of adaptive immunity therefore provides a parsimonious initial trait upon which subsequent adaptations [16] could have gradually introduced additional functionality, leading to systemic regulatory control, refined somatic diversification and long-term memory as observed in contemporary vertebrates. Coincidentally, while the VLRA/VLRB system of jawless vertebrates is functionally similar to the B and T lymphocyte system in jawed vertebrates, no functional equivalent to the MHC (regulatory self/non-self system) has been found in jawless vertebrates [41]. This may support the hypothesis that mucosal adaptive immunity was an ancestral trait shared by both groups, pre-dating distinct transitions to systemic adaptive immunity. In addition, gut-associated lymphoid tissues (GALT) are highly conserved in vertebrates, which further supports the hypothesis of an ancestral adaptive immune system located at the gut interface [89, 119]. Note, no GALT have been reported in jawless fish where lymphoid tissues are located at the tip of the gills [120]. Whether this invalidates the role of the microbiota in the evolution of jawless fish immunity is debatable, since immune control of the microbiota could have originated in pharyngeal tissues at the interface between what later became gut and respiratory tissues in contemporary vertebrates [121, 85].

A crucial assumption of the model is that beneficial microbes were readily available in the external environment, for instance in soils, water and food; and that they may multiply more readily there than potential parasites. Yet, many common beneficial residents of the gut of contemporary vertebrate —such as bacteria from the Bifidobacteria and Lactobacillus groups— do not thrive in the external environment [122, 123]. This might arise from long co-evolution between hosts and select microbial partners, as evidenced by patterns of “phylosymbiosis” [124]; albeit with a strong effect of diet [68]. Indeed, adaptation to living within hosts is usually associated with a marked decrease in genome sizes, preference for anaerobic and warmer environments and specific nutritional requirements such as metabolic pathways specialised in degrading starchy polymers [122, 125]. Long-time microbial partners may be “stuck at the end of the line” [126], such that mechanisms of host control, including diet choice, vertical transmission and metabolic control, might have become sufficient to prevent transitions to parasitism [71]. However, in our model it is not necessary for ‘mutualists’ to be adapted to benefit their hosts yet. Rather, beneficial microbes multiplied slower in the gut of their hosts, relative to parasites. Mutualists may simply replicate metabolic pathways that they use in the external environments and happen to benefit hosts, for instance by fermenting complex sugars or degrading fibres. Arguably, the real challenge presented by host-microbiota interactions might be to prevent “nomadic” [122] and free-living microbes from taking advantage of their host by evolutionary and plastic changes while ensuring the transmission of long-term mutualist partners across host generations and environments. In this context, the model proposed here may be viewed as a representation of the role of immunity in these early associations that pre-date mutual co-evolutionary adaptations between hosts and microbes, highlighting the need for a better understanding of the evolutionary pressures affecting transient microbial partners that have not yet committed to either a fully mutualistic or parasitic lifestyle. Mechanisms of host control such as the targeted sampling of specific food items [127], active recruitment of target microbe species [128] and direct transmission of mutualists from parent to offspring were absent in our minimal model, but may play a key role in the maintenance of host-microbiota cooperation [73]. For simplicity, our model focused a manageably small number of microbe species. However, such mechanisms of recognition-independent host control might simplify the task of immune control down to a comparably small (i.e., 50 − 100 OTUs) number in natural systems, and should make the object of further empirical and theoretical work. Likewise, the extent to which the internal environment of ancestral chordates may have differed from the external environment is unknown, but might be inferred from further research in animals with simple body plans [105].

### 4.4 Conclusion

The present work investigates the potential role of beneficial and antagonistic microbe-host interactions in the ancient evolution of adaptive immunity. Focusing on a minimal model of adaptive immunity, we find the exploitation of a diverse and potentially protective microbiota to be a plausible selective explanation for the appearance and conservation of adaptive immunity. Following this framework, we propose considering stochastic variation in immune receptors and genes —even in the absence of well-identified mechanisms of feedback-mediated clonal reinforcement, for instance alternative splicing of immune proteins in arthropods [129, 16]— as potential candidates for precursory forms of adaptive immunity in non-vertebrate chordates, rather than other traits traditionally viewed as hallmarks of adaptive immunity such as long-term memory and the MHC. In particular, modern comparative approaches to the evolution of adaptive immunity are limited by the lack of species diversity and of ecological variation in jawless fish [130]. Continued research on basal chordates, in which some of these traits may have been overlooked, might inform future theoretical approaches and provide insights into whether our immune system truly evolved to moderate interactions with a diverse and beneficial microbiota.

## 5 Acknowledgements

The authors thank colleagues and collaborators for their feedback in the design and writing process of this work. This work was supported by the University of St Andrews and funded by the DSTL French-UK PhD scheme.

## 6 Data availability

The code used to generate the data and figures in this manuscript can be consulted at https://github.com/lucasmathieuachachera/SDI_coevolution.git.

