## Supplementary material for "Beneficial microbes may have favoured the evolution of adaptive immunity": Methods and Supplementary Figures

### 7 Materials and methods

We simulate the evolution of a population of hosts and a meta-population of microbes, such that each host contains a sub-population of many microbe species. In addition, we model the external environment as a large reservoir of microbes, where constraints on growth rates differ but not microbe-microbe interactions. Microbe evolution is restricted to the mutualism-parasitism continuum and we assume that growth rate is proportional to parasitism, such that selection for increased growth rate is the main driver for increased parasitism.

**Microbe dynamics:** We model each sub-population of microbes as an ecological community using a many-species random generalized Lotka-Volterra model [131]. Microbe time-series within hosts are illustrated in figure 7. For simplicity, microbe-microbe interactions are modelled as pairwise and linear. Within each host and in the external environment in the explicit model, the abundance $X_i$ of microbe species $i$ is controlled following:

$$\frac{dX_i}{dt} = r_i \cdot X_i(t) \cdot (1 - \frac{1}{K} \cdot [\sum_j (\alpha_{ji}) \cdot X_j] - H_i) + \epsilon_i \quad , \tag{1}$$

where $dX_i$ is the incremental change in the abundance of microbe species $i$ at time $t$, $r_i$ is the growth rate of microbe species $i$, $\alpha_{ji}$ is the effect of microbe species $j$ on species $i$, $H_i$ is the sum of host immune effects affecting species $i$ (held to 0 in the external environment) and $K$ is a carrying capacity shared across microbe species. Note $K$ is factorised outside of the sum, but this can be viewed as the more general formula $\frac{dX_i}{dt} = \cdot X_i(t) \cdot (r_i + \sum_j A_{ji} \cdot X_j)$, with $A_{ji} = -r_i \cdot \frac{\alpha_{ji}}{K}$. For the results presented here, $r_i \cdot \alpha_{ji} << K$ which contributes to stabilising ecological dynamics [131]. We used $K_{host} = 100$ in hosts and $K_{env} = 1/2 \cdot n_{hosts} \cdot K_{host}$, such that the environment's carrying capacity was vastly superior to any single host, but sufficiently small to be influenced by exchanges with the host microbiota. As commonly done in random L-V models, we draw $\alpha_{i,j}$ from a normal distribution $\mathcal{N}(\mu_\alpha, \sigma_\alpha)$. When $\alpha_{i,j}$ is negative, microbe species i favours j and when $\alpha_{i,j}$ is positive i competes with j. In figures 3 and 4 we used $\mu_\alpha = 0.7$ and $\sigma_\alpha = 0.3$. As shown in figures 11, and 12 these values produced microbiota leading to near-neutral host fitness, sitting at a saddle point for the frequency of SDI and the species richness of the microbiota, with a high number of different microbiota attractors and high sensitivity to perturbations. For all microbe species, $\alpha_{i,i} = 1$, that is, microbes of the same species compete with one another.

In each assay, microbe interaction terms are sampled randomly as above. Each microbe species is also characterised by its intrinsic growth rate $r_i$ and its effect on the host $\beta_i$. Each microbe →host term $\beta_i$ is initially drawn from a random distribution such that half of all microbe species are beneficial to the host and half are harmful. Note this initial distribution matters very little, because $\beta_i$ were susceptible to rapid evolutionary change. Intrinsic growth rates $r_i$ were calculated using $\beta_i$ such that parasites grew faster than mutualists within hosts, using the following formula. In the external environment, we assumed that $r_i$ and $\beta_i$ were linked by the sigmoid function:

$$r_{i,host} = r_0 - \frac{tanh(\beta_i)}{2} + \xi_{i,host} \quad . \tag{2}$$

In the external environment, we assumed that microbe growth rates were considerably lower, to account for the markedly higher carrying capacity $K_{env}$. This choice also represented the tendency of microbes to grow exponentially faster in the warmer and humid environments characteristic of hosts, compared to external environments. Therefore, microbe growth rates were calculated as:

$$r_{i,env} = \tau \cdot \frac{1}{1+e^{-10\cdot\beta_i}} + \xi_{i,env} \quad , \tag{3}$$

where $\xi_{i,host}$ and $\xi_{i,env}$ are random perturbations drawn each time-step following $\mathcal{U}(0, 0.1)$ and $\mathcal{U}(0, 0.03)$, accounting for stochastic variation in factors uncontrolled by hosts. We bounded $r_i$ to the interval [0,0.5], with $r_0 = 0.2$ guaranteeing higher growth rates within hosts. As illustrated in supplementary materials, figure 6, $r_{i,host}$ is almost equivalent to a bounded linear response to $\beta_i$ over the interval (-0.5,.5). On the other hand, $r_{i,env}$ presents a flat distribution when $\tau << 0$ and a sigmoid distribution with an inflexion point at $\beta_i$=0 when $\tau > 0$ , in order to maximise the selection signal for mutualism for near-neutral microbes in the external environment. When $\tau$¿0 parasites grow on average slower than mutualists in the external environments. In the results presented in figure 2 we varied $\tau$ within [-0.05,0.05] to indirectly modulate the proportion of mutualists and parasites. In other figures, we set $\tau = 0.02$ to give mutualists a very moderate advantage in the external environment.

**Microbe evolution:** Rather than assuming that microbes could evolve by evading their host's immune response, we modelled microbe evolution as variance on the mutualism-parasitism spectrum. To avoid computational costs that would come from modelling individual microbes as agents, we also assumed that within each host, microbe sub-populations evolved under strong selection, weak mutation [101]. That is, each new mutation was either fixated or disappeared before the next mutation appeared. Under these assumptions, every t microbial generations and in each host we draw $k$ microbe species to trial for a new mutation –with $k$ determined by the total number of species and rate of microbe evolution. Unless specified otherwise, we introduced on average 0.5 mutations per time-step and per microbe species in each sub-population. Microbes are trialled for mutation both within each host and in the external environment, such that in each sub-population a microbe species $i$ is trialled for new mutations with a probability proportionately derived from its relative abundance.
Mutations on $\beta_i$ are drawn from a normal distribution $\mathcal{N}(1e-2,\, 1e-1)$. This normal distribution is marginally skewed towards positive values to represent the ability of hosts to bias microbe evolution by imposing stressful conditions, for instance by maintaining a high pH and restricting the metabolic substrates available for microbial growth [73]. Each mutation is trialled by comparing $\frac{dX_i}{dt}$ and $\frac{dX_{i,mut}}{dt}$, using $r_i$ and $r_{i,mut}$ with from the formula described above . If $[\frac{dX_{i,mut}}{dt} - \frac{dX_i}{dt}] > 1e^{-3} \cdot X_i$ , we consider that the mutation rapidly reaches fixation in this sub-population. If the trialled population is undergoing stagnation or currently decreasing (i.e., $\frac{dX_i}{dt} <= 0$) we expect mutations on $\beta$ to be subjected to drift and to reach fixation with probability 0.5.

**Meta-population dynamics:** Each time-step, hosts sample microbes from a subset of $k$ free-living microbe species from the environment. The fraction of microbe species sampled $k$ is drawn from a binomial distribution of size n and probability $p$. We used $p = 0.1$ such that on average a subset of 10% of all microbe species present is drawn each time-step. A species is sampled with probability $p_i = \frac{X_{i,env}}{\sum X_{j,env}}$ and sampling is modelled as a positive perturbation $\delta X_i$ drawn from a normal distribution $\mathcal{N}(\frac{K_{host}}{10\cdot S}, \frac{K_{host}}{20\cdot S})$ where $K_{host}$ is the carrying capacity of the target host and $S$ is the total number of species. Similarly, microbes can migrate from hosts to the environment with the same probability and a perturbation drawn from $\mathcal{N}(\frac{K_{env}}{N\cdot K_{host}}, \frac{K_{env}}{2\cdot N\cdot K_{host}})$, where $N$ is the total number of hosts. For simplicity, we did not model negative migrations ($\delta X_i < 0$). That is, we assumed that the loss of microbes due to migration from one reservoir to another was negligible. Each sub-population of a species $i$ may have slightly different $\beta_{i,host}$ which are impacted by migration from other sub-populations. To account for cases where some microbe species are initially absent (or present in very small density) in a host, we modify the average microbe-host interactions $\beta_{i,host}$ within the host using the average microbe-host interactions from the source $\beta_{i,source}$:

$$\beta_{i,host,t+1} = \frac{\beta_{i,host,t} \cdot X_{i,t} + \beta_{i,source,t} \cdot \delta X_{i,t}}{X_{i,t} + \delta X_{i,t}} \quad , \tag{4}$$

We also tested a scenario where hosts could vertically transmit a fraction of their microbiota when they reproduce. In this case, we used the dynamics described for sampling from the external environment applied to $X_{i,parent}$ and $\beta_{i,parent}$ where *parent* is a reproducing organism. We did not find marked difference in the frequency of SDI in populations with non-zero vertical transmission (figure 9); therefore, the effect of vertical transmission was not reported in the main text.

**Host fitness:** Host absolute fitness $\omega_h$ is calculated as the sum of weighted microbe-host interactions $X_i \cdot \beta_i$ in host $h$ over lifetime, that is:

$$\omega = \int_t \sum_{i,\beta_i>0} f_{benefit}(X_i \cdot \beta_i) + \sum_{i,\ \beta_i<0} f_{harm}(X_i \cdot \beta_i) \quad . \tag{5}$$

Benefits obtained by hosts from mutualists are generally non-linear, for instance due to the functional modularity of the microbiota [76], diminishing returns due to saturation in nutrients [132] and requirements for diversity in metabolic function. Consequently, we defined the dose-response benefit function $f_{benefits}(x) = a \cdot log(1 + x/b)$ where $x$ stands for $X_i \cdot \beta_i$, with $a = 4$ and $b = 3$ respectively controlling the asymptote and slope of the curve. Similarly, we defined the dose-response harm function $f_{harm}(x) = -e^{x/2}$. To assess the effect of the choice of benefit function, we also tested Hill functions $f_{benefit}(x) = \frac{a \cdot x^s}{b^s + x^s}$ where $a = 11$ is the left asymptote, $b = 11.5$ is the incurvation point and $s$ controls the curvature of the sigmoid. Parameters values were chosen such that beneficial microbe-host effects would saturate around $K_{host}/10$ (see supplementary materials, figure 5 for a comparison in function of $x$).

Generations of hosts are overlapping, with a rolling subset $N/4$ of hosts is replaced using parents from a subset $N/2$ of mature hosts, with the last remaining $N/4$ hosts considered as undergoing pre-reproductive development (such that 4 generations of different ages are co-existing at all times). Parents are sampled with repetition using fitness-proportional probability $p = (\omega/\bar{\omega})$ where $\bar{\omega}$ are is the normalized average fitness of the subset of population reproducing. New hosts are produced clonally with mutation and are initially free of microbes, and subsequently sample microbes throughout their lifetime as described in *Meta-population dynamics*.

**Host immunity:** Hosts can influence their gut microbiota using either solely innate immunity or a combination of innate immunity and SDI. Innate immune responses are modelled as:

$$H_{i,innate}(t) = R_i \cdot (1 + \sum_{j,\ \beta_j<0} X_j \cdot \beta_j) \quad , \tag{6}$$

where $H_{i,innate}$ is the sum of innate immune effects (e.g., antimicrobial peptides, phagocytosis, etc) targeted at microbe species $i$, $R_i$ is the reactivity of the host's innate immunity to microbe $i$, and $\sum_{j,\beta_j<0} X_j \cdot \beta_j$ is the total harm caused by the microbiota at time t. Hence, innate immune responses are both constitutive and damage-proportional, following from Hamilton and colleagues (2008); much like constitutive and damage-associated inflammatory responses in vertebrates [134, 135].

When using a combination of innate immunity and SDI, the immune response affecting microbe species $i$ depends on the current population of immune cells (lymphocytes) $L_i$ targeting $i$. Within each host, the density of lymphocyte clones $L_i$ is modelled as a stochastic process using sampled microbes as reinforcement:

$$\frac{dL_i(t)}{dt} = \frac{\sum_j X_j}{\sum_j L_j} \cdot [(1-\gamma) \cdot (r \cdot L_i \cdot R_i \cdot X_i + (1-r) \cdot L_i(t)) + \gamma \cdot \sigma] \quad , \tag{7}$$

where $\frac{\sum_j X_j}{\sum_j L_j}$ scales the intensity of the lymphocyte immune responses to the size of the microbial population to prevent issues in young hosts —much like how the microbiota activates immunity in contemporary vertebrate hosts [25]—, $\gamma$ controls the weight of microbe-independent growth due to mutation/recombination, $\sigma$ is a stochastic perturbation and the lymphocyte replacement rate $r$ controls the rate of turnover in clonal selection. Using this dynamic, when $r$ and $\gamma$ are non-zero the lymphocyte population displays both reinforcement of specific responses to sampled microbes ( $r \cdot L_i \cdot R_i \cdot N_i$) and generation of diversity through stochasticity in the variation and selection process, $\gamma \cdot \sigma$. We used $r = 0.7$ and $\gamma = 0.3$ to allow lymphocyte dynamics to be mainly growth-based, with a significant effect of stochasticity.

The total immune response targeting any microbe, $H_i(t)$ is a combination of innate and SDI contributions:

$$H_i(t) = \mu \cdot \lambda_{innate} H_{i,innate} + (1-\mu) \cdot \lambda_{SDI} L_i \quad , \tag{8}$$

where $\lambda_{innate}$ and $\lambda_{SDI}$ scale the global intensity of each type of immune response and $\delta$ determines whether SDI is present. Hereafter $\mu = 0.5$ when hosts can use a combination of innate immunity and SDI; and 1 otherwise.

**Host evolution:** Host evolution is mainly represented by changes to $R_{i,i\in[1,n]}$, that is, how innate immunity responds to each microbe species. We assume that mutations on innate immunity usually affect multiple host-microbe interactions at once. In other words, we assume that innate immune recognition is coarse/non-specific (e.g. TLR receptors for double-stranded DNA, flagellin etc). We model mutations as additive (positive or negative) incremental changes spread across a randomly sampled subset of $R_i$, such that a positive increment generally increases the strength of innate immune responses to microbe $i$, and inversely for a negative increment. The number of receptors modified in each offspring is drawn from a hurdle model of conditional probability 0.1, mean $S/10$ and standard deviation $S/15$. The amplitude of the incremental change in each receptor is drawn from a normal distribution $\mathcal{N}(1e-1, 3e-2)$. This large mutational variation aimed at accounting for the marked variation in immune-related genes in vertebrates and invertebrates [81] and to allow hosts to keep up with fast-evolving microbes.

We also allowed mutational variation to affect the overall intensity of innate and SDI immune responses $\lambda_{innate}$ and $\lambda_{SDI}$ with mutational noise drawn from $\mathcal{N}(0, 6)$ with $\lambda$ usually taking values between 30 and 200 (see figure 4, panel C). Finally, we let the trait of SDI change following a binary absence/presence Bernoulli trial with a high probability of mutation $p = 1e - 2$, to allow selection rather than constraints on rare mutations to drive the loss and fixation of SDI.p

| Trait | Symbol | Value range | Mutation frequency (per individual and generation) | Amplitude of mutational perturbation |
|---|---|---|---|---|
| General intensity of innate immune response | $\lambda_{innate}$ | $[0, 200]$ | .5 | $\mathcal{N}(0, 6)$ |
| General intensity of SDI immune response | $\lambda_{SDI}$ | $[0, 200]$ | .5 | $\mathcal{N}(0, 6)$ |
| Innate immune reactivity to microbe species $i$ | $R_i$ | $[0, 1]$ | 0.01 | $\mathcal{N}(0.1, 0.03)$ |
| Presence/absence of SDI | $\mu$ | $\{0.5, 1\}$ | 0.01 | – |
| Microbe →host interaction term of microbe species $i$ | $\beta_i$ | $[-0.5, 0.5]$ | $0.2 \cdot \frac{X_i}{X}$ | $\mathcal{N}(0.01, 0.1)$ |
| Specific growth rate of microbe $i$ | $r_i$ | $[0.05, 0.5]$ | – | – |

**Table 1. Summary of host traits (no background) and microbe traits (light grey) evolving in the model.** The mutation frequency and amplitude of mutational perturbations are indicated per-offspring and per-gene. For instance, with $n$ microbe species in the system and as many $R_i$ innate immune genes, the total expected number of mutations per host and per generation is $0.01 \cdot n$. Each model gene except $\mu$ represents a cluster of underlying loci affecting a quantitative trait. Mutational perturbations were therefore drawn from a normal distribution, as commonly assumed for quantitative polygenic traits (e.g. Lande 1976 [136]). For microbes, mutation frequency and amplitude is defined per sub-population (e.g., within each host). Microbe growth rates do not directly mutate, but rather are conditioned to microbe-host interaction terms (see equation (2) and (3) in *Microbe dynamics*).

### 8 Supplementary materials

#### 8.1 Interaction terms in random generalised L-V

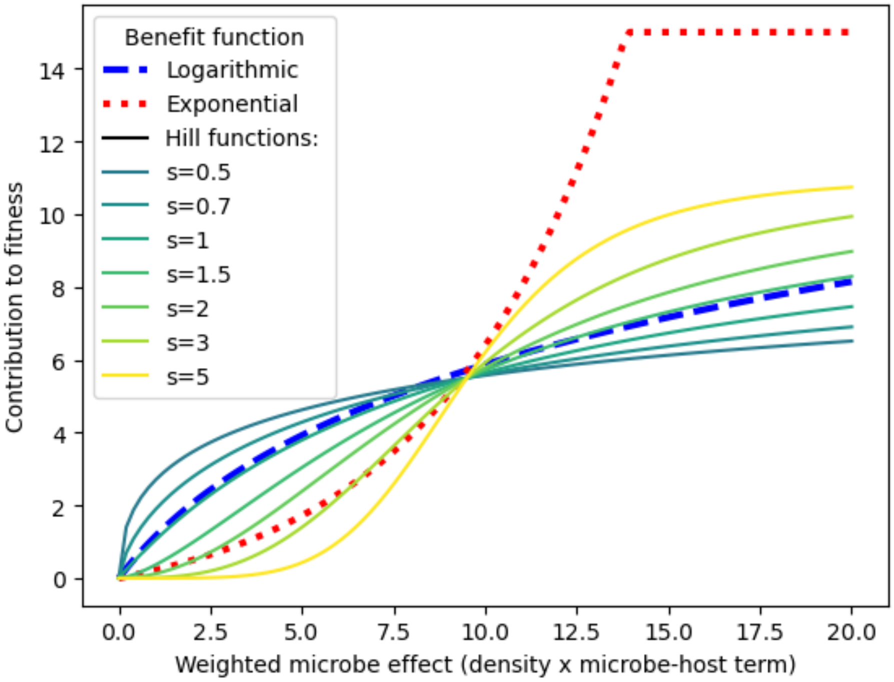


**Figure 5. Comparison of does-response functions for beneficial microbe-host effects**. The functions are defined in Materials and Methods, subsection "Host fitness".

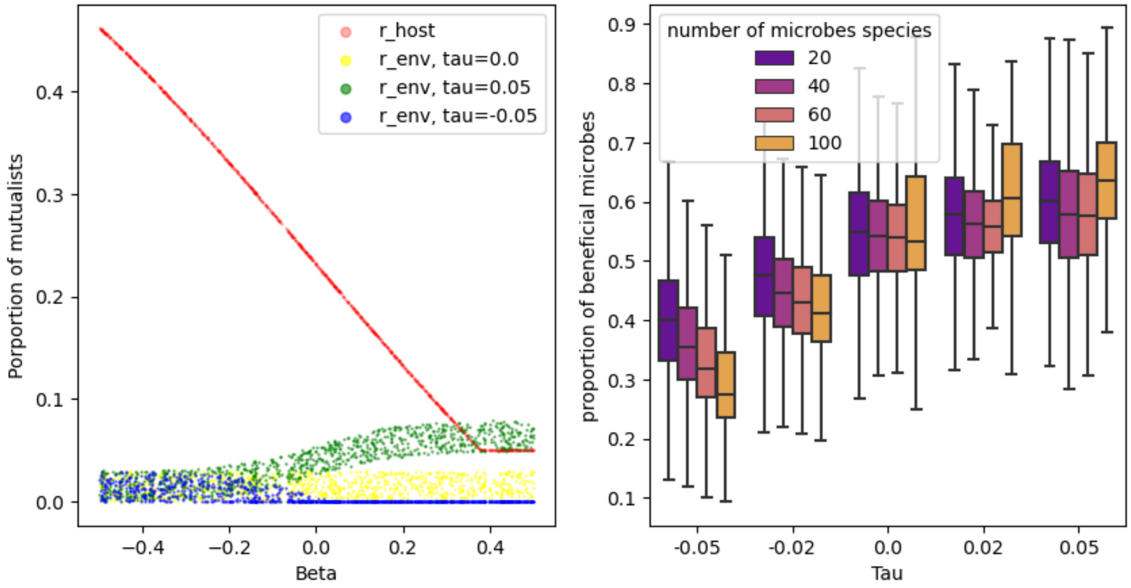


**Figure 6. Effect of $\tau$ on the growth rates (left) and proportion of beneficial microbes (right). Left**: Growth rates within hosts in red are defined as a negative *tanh* function of $\beta$ with a threshold $r_0$ (*tanh* provides a vectorisable near-linear thresholded function). Growth rates in the external environment are defined as Hill functions of $\beta$ and $\tau$, giving a slight advantage in growth/death rates to mutualists ($\tau > 0$) or parasites ($\tau < 0$) in the external environment. **Right**: $\tau$ indirectly controls the proportion of beneficial microbes by constraining the evolution of microbes outside of hosts. Results are shown from populations evolved in the same conditions as figure 2, panel A.

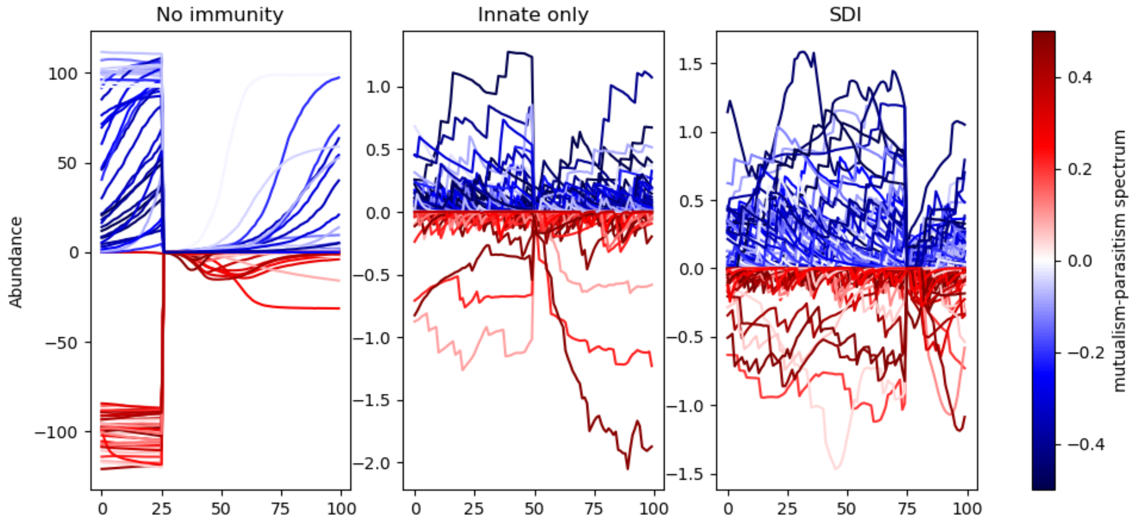


**Figure 7. Time-series of microbes within a host without immune responses (left), with innate immunity (centre) and SDI (right) with $n = 80$.** For better visibility, parasites (in red) are shown with negative abundance. Note the difference in y-axis scale between hosts without immunity and hosts with immunity: microbes proliferate much more in immune-deprived hosts. Compared with SDI, there is higher temporal and inter-microbe variance in abundance.

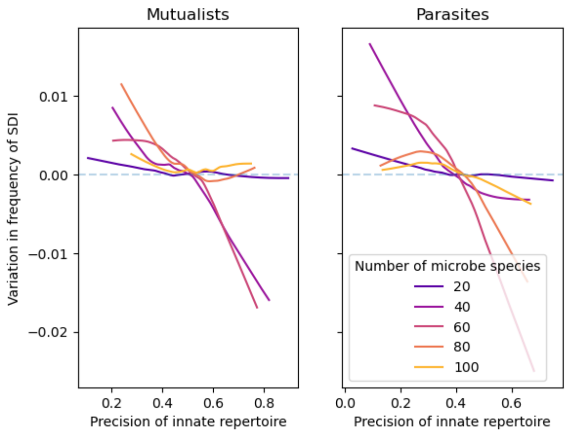


**Figure 8. Effect of innate repertoire performance on variation in SDI frequency.** Repertoire performance is evaluated as the tendency of innate genes $R_i$ to avoid targetting mutualists (left) and target parasites (right). 'Variation in SDI' (y-axis) indicates an increase (positive) or decrease (negative) in the frequency of SDI. Overall, SDI tends to increase in frequency when innate repertoire are performing poorly ($< .5$), particularly when it comes to parasites.

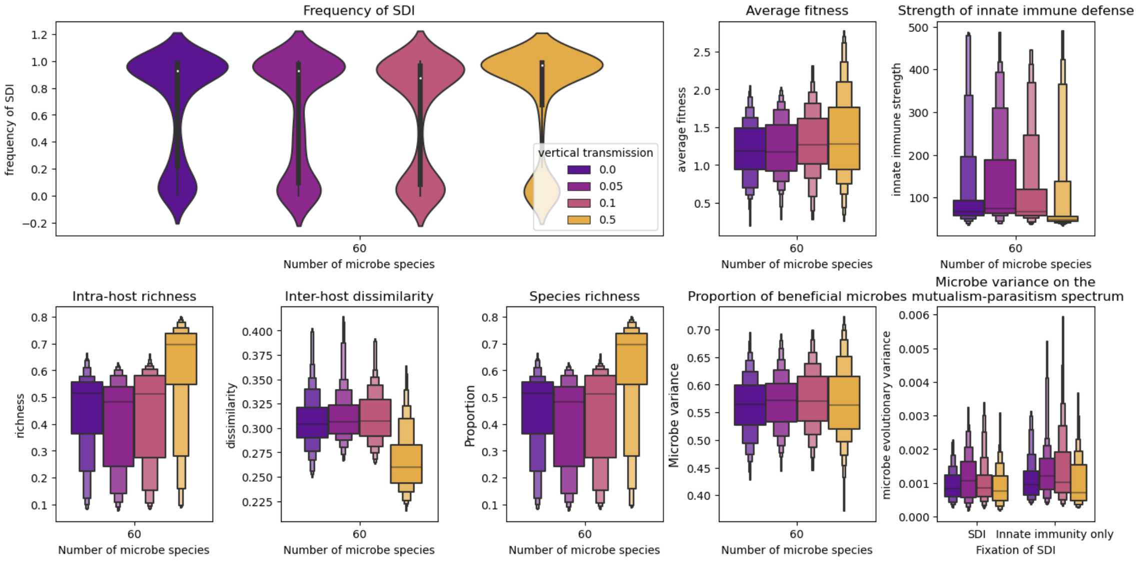


**Figure 9. Sensitivity of the model to vertical transmission of microbes between hosts**. Results are shown with microbe species $n = 60$. Overall, there is little to no effect of vertical transmission on the frequency of SDI. Unsurprisingly, higher rates of vertical transmission (yellow) increase the relative species richness within hosts while decreasing inter-host microbe dissimilarity, by conserving adaptive microbiota across generations.

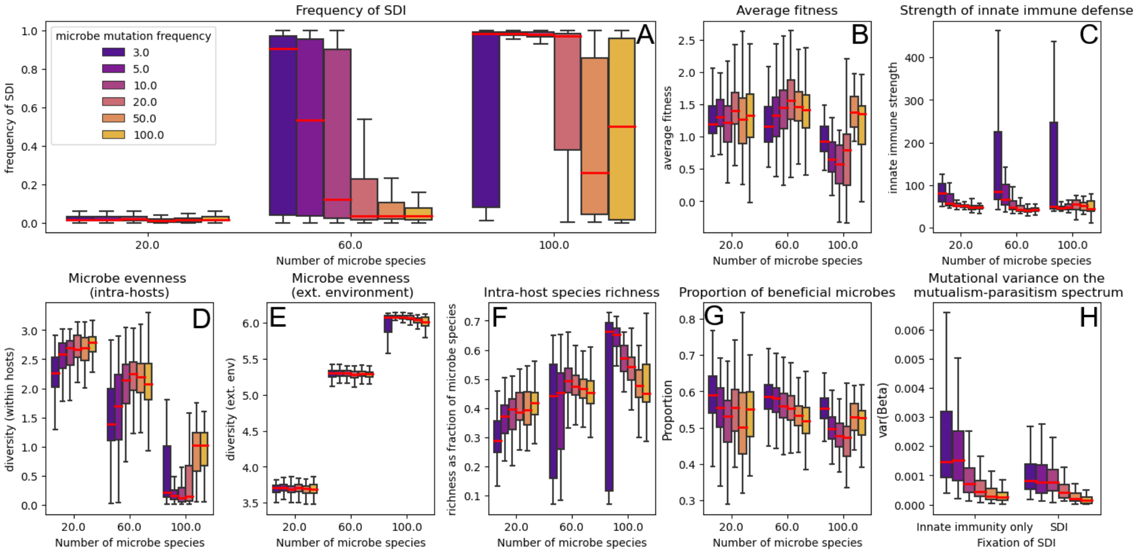


**Figure 10. Effect of microbe mutational frequency on the model**. **A**: At an intermediate number of microbe species ($n = 60$), SDI only evolves with higher mutational frequency. On the other hand, at $n = 100$ SDI is particularly prevalent at intermediate mutational frequencies (5 and 10). Higher microbe mutation rates generally decrease host fitness (B) and increase the intensity of host immune responses (C). Notably, higher microbe mutational variation starkly decreases microbiota evenness within hosts (D) by producing dominant parasites (H).

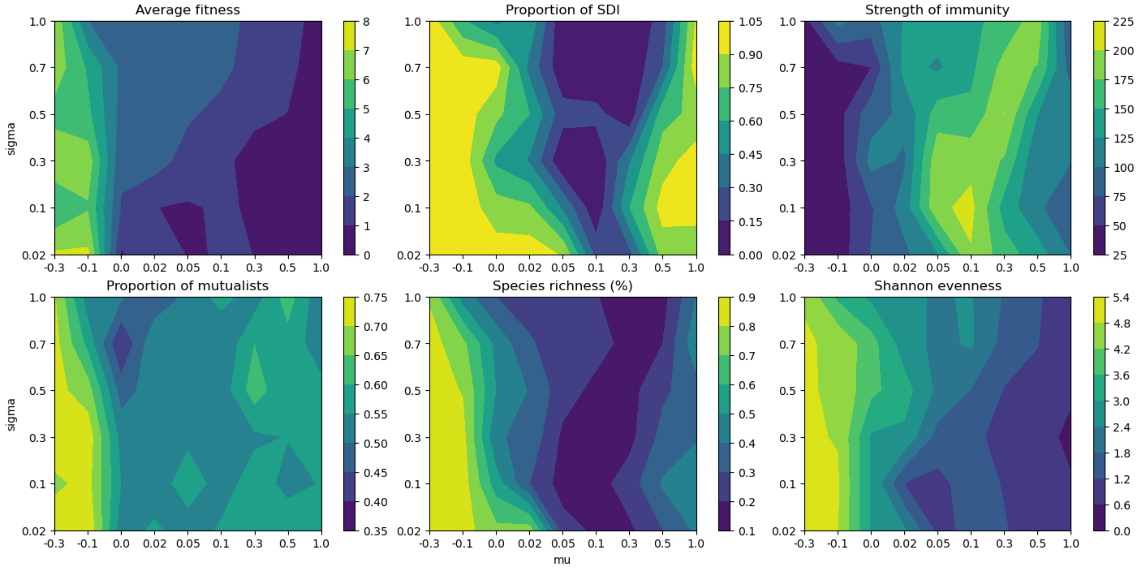


**Figure 11. Sensitivity of host evolution (top row) and microbiota (bottom row) to variation in the mean and variance of normally-distributed linear interaction parameters** $\alpha_{ij}$. Results are shown for a number of microbe species $n = 60$, with $\tau = 0.02$ after 6e4 time-steps of host and microbe evolution (6e3 host generations). Either high competition ($\mu > 0.5$) or high cooperation ($\mu < 0$) favoured SDI in host populations. In accordance with general results on the stability of L-V communities [131], richness (% of non-null abundances) and evenness were markedly higher in populations with weakly competitive or cooperative interactions (bottom row, center and right). Considering that $\mu = .7, \sigma = .3$ was a saddle point for most metrics and that competition is a more common assumption for gut microbes, we used these values in the results presented in this work, unless specified otherwise.

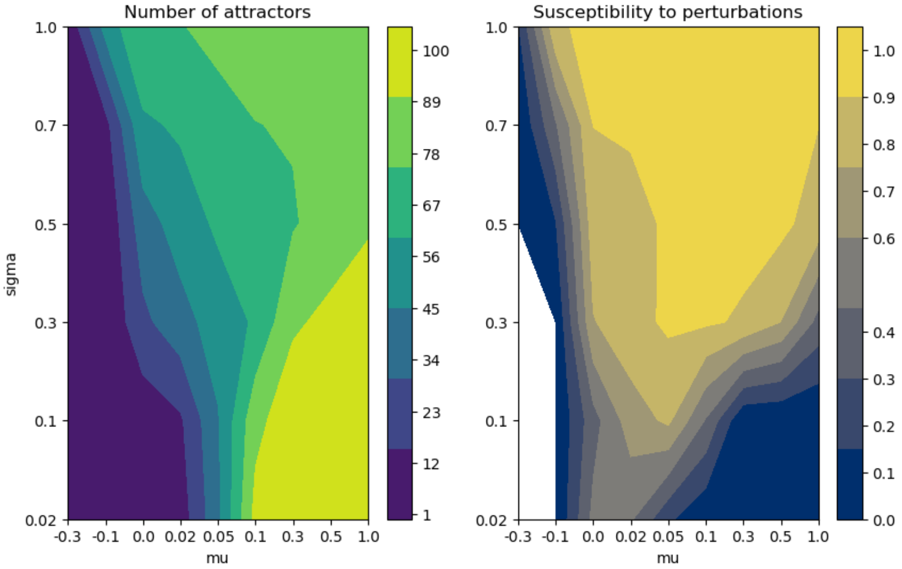


**Figure 12.** Stability of microbe populations in the absence of immune effects. We perturb randomly initialised microbe populations after $t = 100$ time-steps of microbe-microbe interactions in the absence of host immune terms to asses the number and stability of attractors, over 100 trial populations. Left: we use agglomerative clustering to define clusters of microbe abundances. 100 attractors means every set of initial abundances yielded different outcomes; 1 means every microbe population converged towards the same distribution. Right: after perturbing abundances with a normally drawn perturbation of standard deviation $0.3 \cdot \bar{X}$ we assess whether perturbed states still belong in the same cluster as the initial state and count the proportion of perturbations that modify the predicted cluster.
